# Head-to-Head Comparison of Three Methods of Quantifying Competitive Fitness in *C. elegans*

**DOI:** 10.1101/371831

**Authors:** Timothy A. Crombie, Sayran Saber, Ayush Shekhar Saxena, Robyn Egan, Charles F. Baer

**Affiliations:** Department of Biology, University of Florida, Gainesville, FL 32611, USA; Department of Molecular Biosciences, Northwestern University, Evanston, IL 60208, USA; University of Florida Genetics Institute, Gainesville, FL 32611, USA

## Abstract

Organismal fitness is relevant in many contexts in biology. The most meaningful experimental measure of fitness is *competitive* fitness, when two or more entities (e.g., genotypes) are allowed to compete directly. In theory, competitive fitness is simple to measure: an experimental population is initiated with the different types in known proportions and allowed to evolve under experimental conditions to a predefined endpoint. In practice, there are several obstacles to obtaining robust estimates of competitive fitness in multicellular organisms, the most pervasive of which is simply the time it takes to count many individuals of different types from many replicate populations. Methods by which counting can be automated in high throughput are desirable, but for automated methods to be useful, the bias and technical variance associated with the method must be (a) known, and (b) sufficiently small relative to other sources of bias and variance to make the effort worthwhile.

The nematode *Caenorhabditis elegans* is an important model organism, and the fitness effects of genotype and environmental conditions are often of interest. We report a comparison of three experimental methods of quantifying competitive fitness, in which wild-type strains are competed against GFP-marked competitors under standard laboratory conditions. Population samples were split into three replicates and counted (1) “by eye” from a saved image, (2) from the same image using CellProfiler image analysis software, and (3) with a large particle flow cytometer (a “worm sorter”). From 720 replicate samples, neither the frequency of wild-type worms nor the among-sample variance differed significantly between the three methods. CellProfiler and the worm sorter provide <u>at least</u> a tenfold increase in sample handling speed with little (if any) bias or increase in variance.

---

In the context of evolutionary biology, *fitness* is the contribution of an individual to the next generation. Researchers working with *C. elegans* and related nematodes are often interested in comparing the average fitness of different strains. The most straightforward way to quantify fitness is to count the total number of offspring produced by an individual over the course of its lifetime. The number of offspring produced over the lifetime of an individual *i* is its *absolute fitness* (usually depicted *W_i_*). *Relative fitness*, *w_i_*, is the absolute fitness of an individual scaled relative to that of a reference, usually either the most fit individual in the population, i.e, 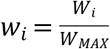 or the population mean, 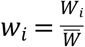.

From the perspective of evolution, only relative fitness matters. All else equal, greater absolute fitness means greater relative fitness. However, all else is often not equal, for several reasons. First, demography matters: offspring produced early in an individual’s life contribute more to fitness than offspring produced late in life [1]. More importantly, however, interactions between individuals can influence fitness in ways that will not be apparent if the different strains are not allowed to interact directly. Also, differences in relative fitness may often only be manifested under competitive conditions [2], because small differences in performance which would have no detectable effect on fecundity (e.g., sprint speed in gazelles) may translate into qualitative differences in fitness (e.g., which gazelle gets caught and eaten by the lion).

A standard laboratory assay of competitive fitness in many organisms, including Caenorhabditis, is to allow different strains of interest (“focal” strains, usually strain=genotype) to compete against a standard, marked competitor strain. Experimental populations are initiated with a known number of focal and competitor individuals, and the population allowed to grow until a pre-defined endpoint, at which time the individuals are either enumerated in totality or sampled. The proportion of individuals of the focal strain at the assay endpoint is designated *p*; the proportion of competitors is 1-*p*. The ratio *p*/*(1-p)*, often called the “competitive index”, *CI*, provides an estimate of the fitness of the focal strain relative to the marked competitor. Relative competitive fitness of strains *i* and *j* can be assessed by comparison of *p_i_*/*p_j_*. There are several indexes of competitive fitness, all of which are fundamentally based on the proportion *p* [3].

Experimental measurement of competitive fitness in many organisms, including Caenorhabditis, has been greatly facilitated by the availability of heritable fluorescent markers, e.g., GFP, which can be scored in much higher throughput than traditional phenotypic markers such as dumpy or unc mutants. The simplest competitive fitness assay is to pick a known number of worms of the focal and fluorescently-marked competitor strains onto a seeded plate, incubate the plate until the food is consumed, then count the worms under transmitted light and fluorescent light. Focal and competitor worms alike will be visible under transmitted light, whereas only the competitor worms will visible under the fluorescent light (Figure 1). Images of the plate can be captured under transmitted and fluorescent light and worms counted, either by eye or by means of image analysis software.

**Figure 1.**
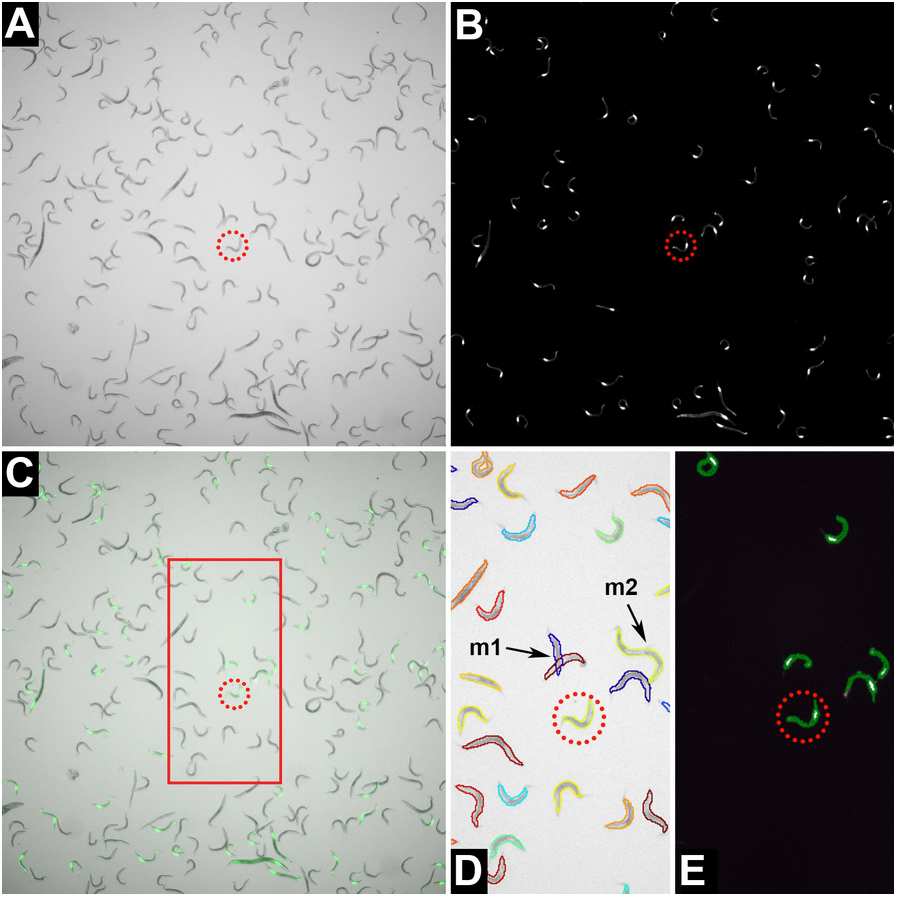
Example of a comparison of the same sample of worms taken under bright-field (A) and under 470 nm fluorescent light (B). The bright-field and fluorescence images are merged showing that the images are exact overlays (C). Worms that appear under fluorescent light are the GFP-marked competitor; worms that do not fluoresce are the focal type. The difference between the number of worms visible in (A) and (B) is the fraction *p* of the focal type. The red circles highlight the same individual in all images. Panels (D) and (E) show CellProfiler generated worm outlines and GFP objects respectively for the area bound by the red rectangle in (C). Occasional CellProfiler worm untangling errors are shown in (D); “m1” shows misaligned worm outlines for the overlapping worms, “m2” shows two worms mistaken as one.

Two methods exist by which the throughput of competitive fitness assays can be significantly increased. First, worms can be washed into wells of a microtiter plate and a motorized stage used to automate the image capture, followed by automated counting by image analysis. Second, a large-particle flow cytometer (aka, a “worm sorter”) can be employed. The latter two methods involve significant initial investment, especially the worm sorter. However, given that the relevant hardware is available, it is useful to know the time/accuracy trade-offs involved with the different methods.

Here we provide a head-to-head comparison of three methods of quantifying competitive fitness in *C. elegans*. Method 1 is our standard “by eye” competitive fitness assay, in which nothing is automated [4]. Method 2 employs the image-analysis software CellProfiler [5,6], applied to data collected from worms washed from agar plates into wells of a 96-well plate. Method 3 involves using a Union Biometrica BioSorter large particle flow cytometer to count worms washed from agar plates into wells of a 96-well plate. Our primary interests are: (1) quantifying the variability of the data, and (2) quantifying the time per datum collected.

## Results

### (i) Variability

Measures of variance are often correlated with the mean, which can confound inferences regarding differences in variance when both quantities differ between groups [7,8]. To begin, we assessed three estimators of relative fitness: the frequency of the focal type, *p*, the competitive index *CI = p/(1-p),* and *log(CI)*. For each estimator of relative fitness we calculated two measures of variation: the within-block standard deviation (SD) of a given focal strain/competitor strain/method combination and the mean within-block Brown-Forsythe statistic 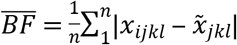, where *x̃_jkl_* is the block median of the estimator of relative fitness of focal strain *j* against competitor strain *k* using method *l*, and *n* is the number of observations in block of a given focal strain/competitor strain/method combination.

Plots of the mean-variance relationships are shown in Figure 2. The correlations are weakest for *log(CI)*, slightly greater for *p* and nearly perfect for *CI* (≈ +1, as expected). For each of the three measures of relative fitness, SD_*log(CI)*_ is less correlated with the mean than is BF_*log(CI)*_. Given these findings, our assessment of the three assay methods is based on SD_*log(CI)*_.

**Figure 2.**
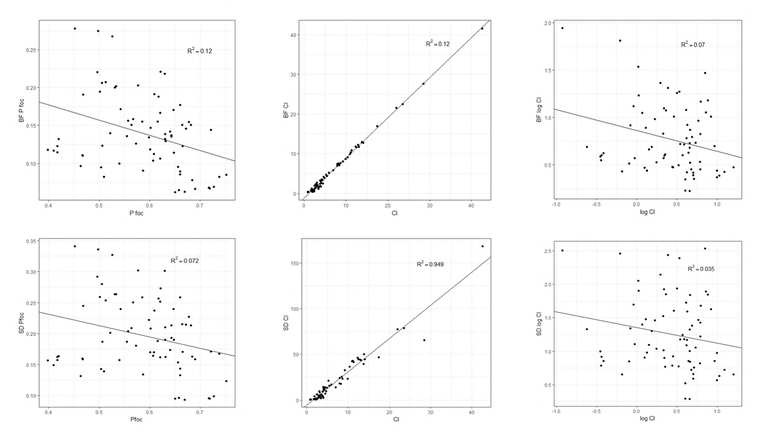
Plots of measures of variability (y-axis) vs. measures of competitive fitness (x-axis). Top panels show the Brown-Forsythe statistic as the measure of variability, Bottom panels show the standard deviation as the measure of variability. Left panels show the frequency of the focal type (“pfoc”) as the measure of competitive fitness, middle panels show CI (= *p*/(1-*p*), right panels show log(CI).

Box-plots of *p* and SD_*log(CI)*_ for the three methods, averaged over focal strains and competitor strains, are shown in Figure 3. The data are summarized in Supplementary Table S1 and raw data are given in Supplementary Table S2. Averaged over focal strains and competitors, *p* does not differ significantly between the three methods (F_2,69.1_=1.71, P>0.18), although it is slightly larger when estimated by CellProfiler. SD_*log(CI)*_ does not differ significantly between the three methods (F_2,49.5_ = 2.23, P>0.12), but CellProfiler is slightly less variable than the other two methods. However, because of the weak negative correlation between mean and variance of *log(CI)*, it is uncertain if CellProfiler is inherently slightly less variable than the other methods, or if the slightly lower variability would disappear in cases when the means were equal.

**Figure 3.**
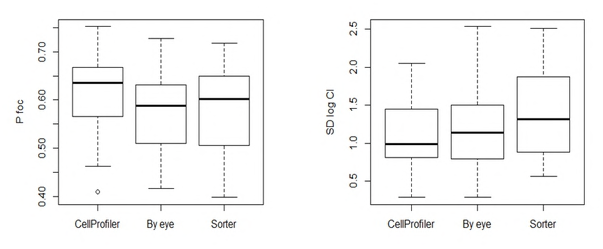
Left panel: boxplots of the frequency of wild-type individuals (“pfoc”) for the three methods. Right panel: standard deviation of log(CI) for the three methods. Each data point is the average of a block/competitor strain/focal strain combination.

The repeatability of the “by eye” and CellProfiler methods can be assessed by counting the same image(s) twice. To quantify the repeatability of the by eye method, the same counter (SS) re-counted a subset of 59 images approximately one month after the original counts. The correlation between the two counts was >99.9% for both the total count and the GFP count. The mean absolute difference between the two counts, expressed as a fraction of the average of the two counts, was 0.73% for the total count and 0.46% for the GFP count. The correlation between the proportion of wild-type worms, *p*, between the two counts is 99.8%. We re-counted all 720 images counted by CellProfiler; the counts were exactly the same in every case.

Unlike images, worm sorter samples are ephemeral. To assess repeatability of the worm sorter counts, we split samples from the same assay well into two wells on the counting plate. The mean absolute difference in the proportion of wild-type worms, *p*, is 4.6%, and the correlation between *p* in the two samples is 0.98. Note, however, that the variation among the two sorter samples includes sampling variance (= technical variance), whereas the variation between the two samples by eye and by CellProfiler only includes counting error.

### (ii) Time per datum

Each of the three methods has unique investments of time associated with it; we return to those in the Discussion. For the head-to-head comparison, each sample was divided into three aliquots (see Methods), thus the clock starts at the time the samples were divided. The “by eye” and CellProfiler methods both involve the analysis of images captured from samples in a 96-well plate. It took approximately 20 minutes to capture the 96(x2) images from one 96-well plate with the stage movement rate set to “slow” (faster image capture is possible), or about 12.5 seconds/sample.

An experienced counter (SS) took ~5-7 minutes to count both images for one sample by eye (bright field and GFP; *n̅*_*BF*_≈176, *n̅*_*GFP*_≈77). The rate at which images can be processed by CellProfiler will depend on the computer platform as well as the images. Working with a far-from-state-of-the-art Dell Optiplex 9020 PC running Windows 7, CellProfiler averaged 19.5 ± 8 minutes to process a 96-well plate, or about 12.5 seconds per sample.

As we applied it, the worm sorter took about 50 minutes to collect data from a 96-well plate, or about 30 seconds per sample. Once data are acquired by the sorter, counts can be obtained computationally essentially instantaneously.

## Discussion

The results of this study are good news for worm biologists wanting reliable high-throughput estimates of competitive fitness using fluorescently-labeled strains. Automated image-capture with CellProfiler and the BioSorter both provide estimates of competitive fitness that are nearly unbiased and no more inherently variable than the human eye. For samples of a few hundred worms, CellProfiler and the Biosorter can count worms at a rate that is approximately an order of magnitude faster than even a well-trained, well-motivated human, and the time-savings will increase as sample size increases.

A caveat is that both CellProfiler and the Biosorter require a non-trivial initial investment of time to optimize the data-collection protocol, whereas a sighted human can begin collecting data right away. In our experience the CellProfiler image analysis pipelines took nearly 40-man hours to develop and implement. However, less time will be required to optimize our existing pipelines (Supplementary File S3) for a different imaging system.

## Methods and Materials

### Strains

Three *C. elegans* strains (N2, PB306, and CB4856) were selected as focal strains for the competitive fitness assays. The competitor strains ST2 and VP604 were chosen based on bright, constitutive expression of GFP reporter alleles in all developmental stages. ST2 *ncIs2* [*pH20::GFP +* pBlueScript] expresses GFP in nearly all neurons and VP604 *kbIs24* [*Pgpdh-1::dsRed2;Pmyo-2:: GFP;unc-119*] X expresses GFP robustly in the pharynx.

### Competitive fitness assay

Five replicate blocks of the competitive fitness assay were conducted in the spring of 2016. Blocks consisted of 24 replicate competitions for all 18 combinations of focal and competitor strains and assay methods, except in one block for which “by eye” data were not collected. Blocks were initiated by bleach-synchronization of strains that had been maintained as mix-staged populations on standard 100 mm NGM plates. The competition between strains began by transferring a single L4-stage focal hermaphrodite and a single L4-stage, GFP-marked competitor hermaphrodite to the same well of a 24-well plate. Each 24-well plate contained NGMA agar supplemented with nystatin and streptomycin and seeded with 10 μl of OP50-1 *E. coli*. Replicates for each of the six competitions were distributed equally among six 24-well plates, i.e. four replicates per 24-well plate. Competition between the focal and competitor strains persisted for 168 h at 20 °C at which point the food source in nearly all wells had been completely consumed. The resulting nematode populations were then washed from the competition plates by adding 1.5 ml of M9 buffer to each well with a repeat pipettor, aspirating up and down three times with a disposable transfer pipet, then transferring the nematode suspension to a 2 ml capacity 96-well plate. Once filled, the 96-well plate was centrifuged for 1 min at 1,000 g, and the supernatant was aspirated with an 8-channel strip aspirator. In order to retain the nematode pellet, the strip aspirator was set to leave 100 μl in each well. The wells in the 96-well plate were then washed once more by adding 1.4 ml of M9 buffer, centrifuging, and aspirating, followed by the addition of 1.4 ml of M9. A 12-channel pipet was then used to gently mix the nematode suspension by pipetting up and down five times. Once mixed, a 110 μl sample of the resulting worm suspension was transferred into the appropriate rows of two 96-well, clear bottom plates. One of the 96-well plates was used to quantify competitive fitness using the image-based methods (manual counting and CellProfiler counting) while the other was used for sorter-based counting.

### Imaging

Quantification of competitive fitness by image-based methods requires both a bright-field and paired GFP fluorescence image to identify focal animals (non-fluorescent) from competitors (fluorescent). To ensure that nematodes do not move between the bright-field and GFP fluorescence images, levamisole was added to the wells at a final concentration of 5 mM. Bright-field and GFP fluorescence images (470±20 nm excitation, 525±25 nm emission) were captured at 20x or 30x magnification for each well using an automated epifluorescence microscope (IX-70, Olympus, Pittsburgh, PA, USA) fitted with a CCD camera (Retiga-2000R, QImaging, Surrey, BC, Canada), XYZ stage and focus motors (Prior Scientific, Cambridge, UK), and controlled by Image Pro-plus ^®^ software (Media Cybernetics, Rockville, MD, USA). The bright-field and GFP fluorescence image pairs for each well were saved as a single stacked image before being used for the image-based competitive fitness quantification methods.

### Quantification method

#### (1) “By eye”

*Manual* counts of focal and competitor worms were obtained as follows. Stacked images consisting of a bright-field slice and GFP fluorescence slice were imported into ImageJ software (http://rsb.info.nih.gov/ij/). A rectangle encompassing approximately 200 worms was superimposed on the bright field image slice and the worms within the rectangle counted (Figure 1a). We used the multipoint tool in ImageJ to facilitate counting worms. The multipoint tool allows the user to mark each worm on the image as it is counted and therefore reduces the chances of miscounting worms. The same rectangle was then superimposed on the GFP image slice and the fluorescent worms visible in that image counted using a different counter type in the multipoint tool (Figure 1b). The number of focal and competitor worms within a given well were then exported from the ImageJ results window and the frequency of the focal type, *p*, the competitive index *CI = p/(1-p),* and *log(CI)* were calculated in excel.

#### (2) CellProfiler

We developed an image analysis pipeline using CellProfiler software to automatically quantify competitive fitness in a given well using the paired bright-field and GFP fluorescence images (Supplementary File S3). CellProfiler is a free, open-source software developed to automate quantitative measurement of phenotypes from large collections of images [6]. Implementation of the Worm Toolbox modules within CellProfiler allows for the automated detection of individual nematodes from bright-filed images despite instances of nematodes clustering or overlapping [5]. Further processing of the paired GFP fluorescence images allows our pipeline to distinguish between focal (non-fluorescent) or competitor (fluorescent) phenotypes, and thus calculate all three estimators of relative fitness in a given image, i.e., frequency of the focal type, *p*, the competitive index *CI = p/(1-p),* and *log(CI)*. Quantification of relative fitness with CellProfiler begins by building a probabilistic model of nematode shape hereafter referred to as a “worm model”. The first step in generating a worm model is to manually identify examples of single, non-overlapping nematodes using the UntangleWorms module from Worm Toolbox configured in the “train” mode. We used the “identifying_single_worms_for_worm_model.cpproj” project file to identify ≥ 50 single, non-overlapping nematodes from 5-10 random bright-field images within a particular block. The resulting “singleworm.png” binary output images were then used as input images for the “create_new_worm_model.cpproj” project file, which processes the single worms in the images to generate the worm model for a particular block. We generated a different worm model for each block due to variations in image magnification and nematode shape across blocks. We then used the “UntangleWormsCountGFP_[Assay_#].cpproj” project files to quantifying the number of focal and competitor nematodes in all images for each block. These project files each use their own image processing pipeline and block-specific worm model. The image processing pipeline for blocks 10 – 13 include crop modules, while block 8 does not. The cropping is included to help correct for extremely uneven illumination in bright-field images, wherein well edges appear dark and centers appear bright. Image processing pipelines for all blocks include modules to detect and correct for any uneven illumination that remains. Subsequent modules then identify putative nematode objects from the background via thresholding. Next, individual nematode outlines are identified from clusters of nematodes or debris using the block specific worm model. Using the paired GFP fluorescence image, fluorescent objects in each nematode outline are identified and counted by thresholding. The pipelines output a .csv file containing the number of nematode objects in a well, along with the number of children GFP objects counted within each nematode object. A particular nematode is assigned a focal genotype if it contains no GFP objects, or a competitor genotype if it contains one or more GFP objects (Figure 1d). Finally, the frequency of the focal type, *p*, the competitive index *CI = p/(1-p),* and *log(CI)* are calculated for each well.

#### (3) BioSorter

Our BioSorter competitive fitness analysis pipeline is presented in detail in Supplementary Appendix A1 of [4]; we reprise the basics here. After sample preparation, we aliquot 110 μl of the sample into a well of a 96-well plate, which the LP Sampler aspirates to the sorter. The LP Sampler script is optimized for 100 μl input, but there is a risk of incorporating bubbles in the fluid line if the sample is short. We include the extra 10 μl for redundancy, and four wash steps with the LP Sampler to ensure we pick up most of the sample. In a typical wash step, the LP Sampler aspirates and dispenses water, cumulatively picking up most of the sample over the course of four wash steps. Potentially leftover samples are unlikely to introduce bias in a competitive assay.

The competitive fitness assay was extended until plates were starved of *E. coli* food (or nearly so), which greatly reduces the frequency of false positives in the sorter counts. *E. coli* tends to clump together and register as many small events (particles), decreasing the signal to noise ratio for smaller worms.

### Data analysis

The dependent variable for purposes of quantifying variation is the standard deviation of log(CI), SD_*log(CI)*_. CI was measured from 24 replicate plates from each combination of focal strain (N2, PB306, CB4856), competitor strain (ST2, VP604), and quantification method (by eye, CellProfiler, Sorter) in five assay blocks. Thus, there are five estimates of SD_*log(CI)*_ for each unique combination of focal strain/competitor strain/method (four estimates for “by eye”). Method is the independent variable of interest; for the purposes of this study we are not interested in the variation among focal strains or competitor strains. However, because the sample sizes are small, we treat focal strain, competitor strain, and their interactions as fixed effects. Data were analyzed by general linear model (GLM) as implemented in the MIXED procedure of SAS v. 9.4. Variance components were estimated by restricted maximum likelihood (REML). The full linear model is 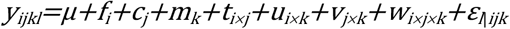, where *y_ijkl_* is the estimate of SD_*log(CI)*_ in block *l*, *μ* is the overall mean, *f_i_* is the effect of focal strain *i*, *c_j_* is the effect of competitor strain *j*, *m_k_* is the effect of method *k*, *t_i×j_* is the effect of the interaction between focal strain *i* and competitor strain *j*, *u_i×k_* is the effect of the interaction between focal strain *i* and method *k*, *v_j×k_* is the interaction between competitor strain *j* and method *k*, *w_i×j×k_* is the effect of the three-way interaction, and 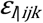 is residual (among-block) variance. We initially estimated the residual variance separately for each focal/competitor/method combination, then pooled the residual variance over different combinations of groups, using the minimum corrected Akaike’s Information Criterion (AICc) as the criterion for the best model. Similarly, competitor strain, focal strain, and their interactions were removed and the AICc calculated. The smallest AICc was given by the model with only method included as a fixed effect and the residual variance estimated separately for each method, pooling residual variance over focal and competitor strains within a method. Significance of fixed effects was assessed by F-test on type III sums of squares.

We repeated the above analysis for the fraction of focal worms, *p*, with block included as an additional random effect and replicate (nested within block) as the unit of observation. The block for which we did not collect “by eye” data was omitted from the analysis.

## Acknowledgments

We thank Joanna Dembek and Asher Shoucair for assistance in the lab. Support was provided by NIH grants R01GM107227 to CFB and E. C. Andersen, and S1010OD012006 to CFB. SS was supported by a graduate fellowship from the Higher Committee for Education Development in Iraq.

## Supplementary Material

**Supplementary Table S1.** An Excel spreadsheet containing summary statistics.

**Supplementary Table S2.** An Excel spreadsheet containing the raw data.

**Supplementary File S3:** This zip file contains sample bright-field and GFP fluorescence image pairs as well as CellProfiler project files. CellProfiler project files 1-2 are used to train and create worm models. CellProfiler project file 3 is used to score focal and competitor worms in image pairs and output scores as .csv files.

